# Preliminary study on the application of PspA as carrier

**DOI:** 10.1101/305102

**Authors:** Lichan-Wang, Yajun-Tan, Chen-Wei, Huajie-Zhang, Peng-Luo, Shumin-Zhang, Xiao-Ma

## Abstract

The aim of the study is to research the feasibility of pneumococcal surface protein A (PspA) using as carrier protein. Three recombinant pneumococcal surface proteins A (come from family1 and family 2) were expressed by prokaryotic expression system and were conjugated to group A meningococcal polysaccharide (GAMP) to make three polysaccharide-protein conjugates. The conjugates, un-conjugated proteins, GAMP and GAMP-TT vaccine bulk (used as positive control) were immunized to mice and their immune effects were evaluated by the method of ELISA, FCM and SBA. The results showed that the polysaccharide-protein conjugates can produce higher levels of anti-GAMP IgG titers (P < 0.05), higher ratios of Th1/Th2 (P < 0.05) and higher levels of serum bactericidal activity (P < 0.05) compared with the un-conjugated GAMP. The conjugation of PspAs to GAMP also enhanced the anti-PspA responses compared with un-conjugated PspAs except PspA3. In conclusion, all the results indicated that three PspAs were suitable carrier protein as demonstrated by the characteristics of a T-cell dependent response to the GAMP, and would protect against group A of epidemic cerebrospinal meningitis and also have the potential to provide broad protection from Streptococcus pneumonia.

## Introduction

Invasive bacteria, such as *Haemophilus influenza type b* (Hib), *Neisseria meningitides* and *Streptococcus pneumonia*, the most important antigen is polysaccharide on the surface of them. Capsular polysaccharide is thymus independent antigen; immunization of infants and young children with these antigens does not induce well and long-lasting protective levels of serum antibodies. The success of Hib glycol-conjugate vaccine highlighted the advantages of converting polysaccharides into T-dependent antigens by chemical conjugation to carrier proteins. Tetanus toxoid, diphtheria toxoid and the non-toxic mutant of diphtheria toxin, CRM197 are commonly used as protein carrier. With the increasing of polysaccharide conjugate vaccines, the development of new protein carrier is necessary for avoiding immune tolerance, immune interference or immune suppression due to repeated using the same carrier (1,2). In addition, the imperfection of the polysaccharide and conjugate vaccine is that the protection of them can not cover all serotypes of the bacteria strains. Serotype conversion limits their development and decreases their immune protecting coverage. Thus increasing the selection of carrier protein or developing the new conjugate vaccine without serotype limitation is important and necessary in future.

In order to solve the above problems, we chose pneumococcal surface protein A (PspA) as research subject. PspA is virulence associated protein found on the surface of all clinical isolates of *S. pneumonia* (3) and interferes with opsonophagocytosis by interfering with complement deposition on the bacterial surface (4,5). It has five domains: a single peptide, α-helical and charged N-terminal domain, a proline-rich region, a choline-binding domain and a short hydrophobic tail. PspA is relatively variable at the DNA and protein sequence levels. According to the sequences of α-helical region, PspAs have been divided into six clades, which belong to three families (6, 7). PspA family 1 contains two clades (1 and 2), PspA family 2 contains three clades (3, 4 and 5) and PspA family 3 contains one clade (clade 6). Families 1 and 2 are expressed in more than 90 percent of strains (8). Antibodies from different clades of the same family have relatively high cross reactivity and cross protection, but lower about clades from different families. Based on its structural diversity, it has been suggested that PspA-based vaccine should contain at least one clade from each of the two major families in order to elicit broad protection (9).

In our study, different clades of recombinant PspA protein (clade1, clade2 and clade3) were prepared and conjugated to group A meningococcal polysaccharide (GAMP) as carrier protein to make available a compounds, which not only prevent epidemic cerebrospinal meningitis but also has the potential to provide protection from Streptococcus pneumonia. The immune effects, including humoral and cellular immune responses were assessed after being immune to mice. In this article, we focused on the ability of the conjugates to prevent meningococcal infection, and pneumococcal infection will be covered in a follow-up study.

## 2. Materials and methods

### 2.1. Preparation of recombinant PspA proteins

*PspA* gene sequences of bacteria strain DBL6A (GenBank AF071805.1), RX1 (GenBank U89711.1) and EF3296 (GenBank AF071816.1) were obtained from GenBank and synthesized after being codon optimization in accordance with *Escherichia coli* codon preference so as to enhance their protein expression in prokaryotic expression system. The synthesized DNAs were 1143bp for PspA/DBL6A, 1122bp for PspA/ RX1 and 1431bp for PspA/ EF3296, which belong to clade 1, clade 2 and clade 3 respectively. Signal peptide sequences of these DNAs have been excluded. The constructed genes were cloned into plasmid pET-30a (+) vector with Nde I and Xho ı restriction enzyme cutting site. PCR and enzyme digestion were used to identify the correction of linkage. The expression plasmid was transformed into competent *E.coli* BL21 (DE3) and expressed by induction with isopropyl-D-1-thiogalactopyranoside (IPTG). After being purified by several steps of chromatography (GE healthcare), three recombination proteins were obtained, analyzed and identified by SDS-PAGE and Western blot. The following recombinant PspA molecules without His tag were produced, PspA/ DBL6A (named PspA1) which contains 375 amino acids, PspA/ RX1(named PspA2) which contains 369 amino acids, and PspA/ EF3296 (named PspA3) which contains 471 amino acids.

### 2.2 Preparation of conjugates

Cyanogen bromide activation method was used to conjugate three PspA proteins to GAMP respectively, which completed in Beijing Lvzhu Biopharmaceutical Co. Ltd. The procedure briefly contains activation of GAMP with cyanogen bromide, conjugation of polysaccharide derivative with recombinant PspA by carbodiimide method and purification of the compound with SephacrylS-400 column chromatography. The ratio between protein PspA and GAMP was determined by improved Lowry method and Phosphorus assay respectively (10). Table 1 is the results of the conjugates. GAMP-TT (used as positive control) is vaccine bulk provided by division of respiratory tract bacterial vaccine of NIFDC.

**Table1.** The ratios of polysaccharide to protein for GAMP-PspA conjugates

| Groups | Polysaccharide content<br>($\mu\text{g/ml}$) | Protein content<br>($\mu\text{g/ml}$) | Ratio of polysaccharide<br>/ protein |
| --- | --- | --- | --- |
| GAMP-PspA1 | 57.3 | 91.4 | 0.63 |
| GAMP-PspA2 | 63.1 | 100.8 | 0.63 |
| GAMP-PspA3 | 121.5 | 195.6 | 0.62 |

### 2.3 Animal immunization

Groups of Blab/C female mice weighting 16-18g were injected subcutaneously with each of the conjugate preparations or unconjugated controls. The dose of polysaccharide per injection was fixed at 2.5μg. The dose of protein varied from conjugate to conjugate and is presented in table 1. Unconjugated proteins as control were injected 5μg per mice. Mice received 3 doses at day0, day14 and day28 and were bled by retro orbital puncture at day13, day27, day35 and day180. Serum were separated and stored at −20°c for testing. All animal experiments were accordance with Regulations on Management of Laboratory Animal (Ministry of Science and Technology) 1988, China.

### 2.4 Antibody detection

Anti-PspA and anti-GAMP IgG titers at day13, day27, day35 and day180 were assayed respectively by enzyme-linked immune sorbent (ELISA). 3μg/ml PspA and 3μg/ml GAMP were coated in 96-well (Nunc) plates. Sera were 2 fold serially diluted across the plates with the predetermined initial dilution. HRP-conjugated goat anti-mouse IgG (Santa Cruz) and OPD (Ameresco, USA) substrate were used as the second antibody and color agent. The reaction was stopped by adding 50μl of 1 M H2SO4 and was read at 490 nm with microplate reader.

### 2.5 Th1/Th2 detection by FCM

Seven days after the third immunization (days 35), 1ml spleen cells (2×10^6^/mL) from mice of each group were co-cultured with 50ng/ml PMA(sigma), 1μg/ml ionomycin (Sigma) and 10μg/ml BFA(Biolegend) in their final concentration for 4-6 hours at 37 °C with 5%CO_2_ incubator. Washing cells with cell staining buffer and labeling with mouse CD3-PerCP, CD4-FITC antibody (Biolegend) for 20mins in the dark. Then after the cells were fixed with fixation buffer and permeabilization with permeabilization wash buffer, which were labeled with mouse IFN- γ PE and IL-4APC antibody (Biolegend) for 30mins in the dark and then were washed and suspended in cell staining buffer for FCM analysis.

### 2.6 Bactericidal detection

Serum bactericidal assay (SBA) is one of the best methods on assessing the potency of meningococcal polysaccharide vaccine, which mainly detect the functional antibodies contained in serum. Meningococcal strain and baby rabbit complement were provided by Lanzhou institute of biological products. Serum samples and rabbit complement for control were heat inactivated for 30mins at 56˚C for use in the assay. Make serum sample and control sample to 3 fold serial dilutions for a total of 8 dilutions in 96 well U micro titre plates with initial 4 fold dilution. Add 10μl of group A meningococcal strain to each well and gently tap plates to mix for 30mins, then add 10μl of complement or heat inactivated complement to control well and sample well respectively, which cultured for 1hr at 37 °C with 5%CO_2_ incubator. Mix plate and take 10μl from eight serial diluted well onto the THYA plate and tilt plate immediately, make the liquid flow into one, about 2-3 cm. Other samples and so on. The plates were placed in room temperature for 20mins and transfer to 37°C with 5%CO_2_ incubator for overnight. After overnight incubation, THYA plates were covered with TTC superstardom agar and using automatic colony counter count colony number after agar solidification. According to the control panel of colony, counting experimental group 50% sterilization drops.

### 2.7 Statistical analysis

IBM SPSS Statistics 20 was used to analysis. The significance of differences in different groups was assessed using a one-way analysis of variance (ANOVA). Pearson correlation was used to analyze the correlation between the results of SBA and antibodies. Kolmogorov-Smirnov was used to test the normality of data. For all comparisons, a P-value of <0.05 was considered to represent statistical significance.

## 3. Results

### 3.1 Production and purification of PspAs

The amplified products were 1000 to 2500 base pairs (bp) by PCR and enzyme digestion using expression plasmids as template, which are consistent with the expected size (figure1 A to B). The three proteins were soluble expressed by Escherichia coli induced with IPTG(final concentration 1mmol/L) at 37 °C for 4hrs, which about 30% to 40% of the amount of total expression. Through three steps of chromatography, the purity of them reached 90 percent (figure1 D to F). Western bot results (figure1 C) showed that the three proteins can react with human serum of clinically diagnosed pneumonia, which indicated the proteins have good antigenicity.

**figure1.**
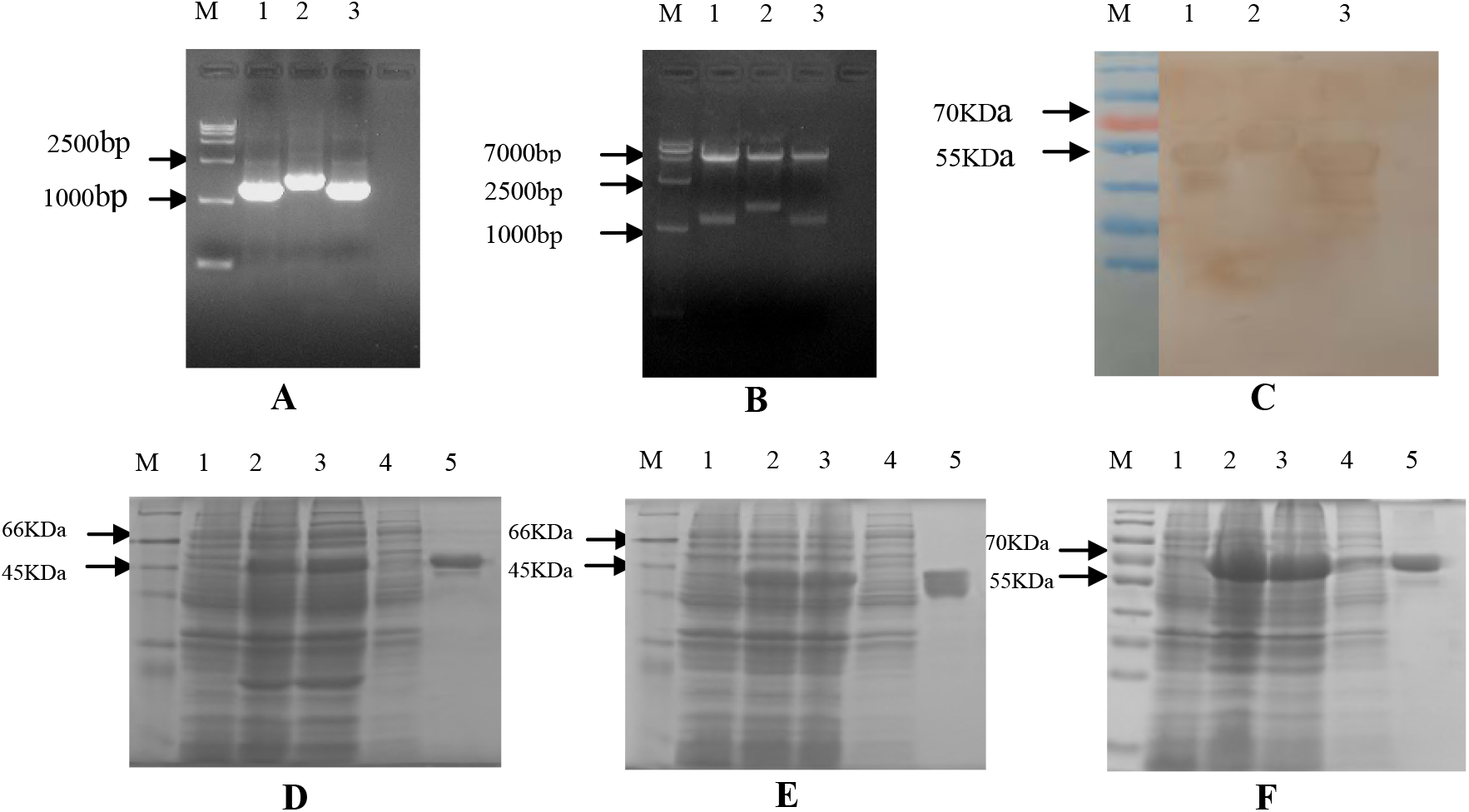
Preparation and identification of three PspA proteins. A to B: Identification results by PCR (A) and enzyme digestion (B) of three PspA expression plasmids. M: DNA Marker, Lane1: PspA1 protein, Lane2: PspA3 protein, Lane3: PspA2 protein. D to F: SDS-PAGE results for recombinant protein PspA1, PspA2 and PspA3. M: Marker; Lane1: empty vector control; Lane2: recombinant protein 4h after induction with IPTG; Lane3: supernatant of recombinant protein after sonication; Lane4: precipitation of recombinant protein after sonication; Lane5: supernatant of recombinant protein after purification. C: Western blot results of PspA proteins. M: Marker; Lane1: recombinant protein PspA1; Lane2: recombinant protein PspA3; Lane3: recombinant protein PspA2.

### 3.2 Chemical and physical analysis of the conjugates

Three conjugates were synthesized: PspA1-GAMP, PspA2-GAMP and PspA3-GAMP, prepared with cyanogen bromide activation method. The content of protein and polysaccharide were tested by the methods described in Chinese pharmacopoeia (10). The ratio of polysaccharide and protein were calculated according to their concentration. Table 1 shows the results.

### 3.3 Immunogenicity of the conjugates and un-conjugates

The anti-GAMP, anti-PspA and anti-TT titers were calculated through compared with that of PBS control mice sera, and expressed as the reciprocal of geometric mean titers (GMT). The anti-GAMP responses are presented in figure 2. All conjugates induced anti-GAMP responses after one dose, and increased with the second and third booster immunization, which were significantly higher than that of unconjugated GAMP, P < 0.05. The anti-PspA (three clades) and anti-TT responses are presented in table 2. The dose of PspA was almost same for each conjugate due to the ratio of GAMP and PspA is similar. The anti-PspA1 and anti-PspA2 responses following the first dose of conjugate were low; responses were all boosted after the second and third dose, which were significantly higher than that of unconjugated PspA1 and PspA2, P < 0.05. The anti-PspA3 titer to conjugates with PspA3 carrier protein was significantly lower after the third dose than that of unconjugated PspA3, P < 0.05. For GAMP-TT conjugate, anti-TT titer was almost dozens of times lower compared to that of unconjugated TT either after the first immunization or after the booster immunizations, which indicates the conjugates inhibited the immunogenicity of TT.

**Figure2.**
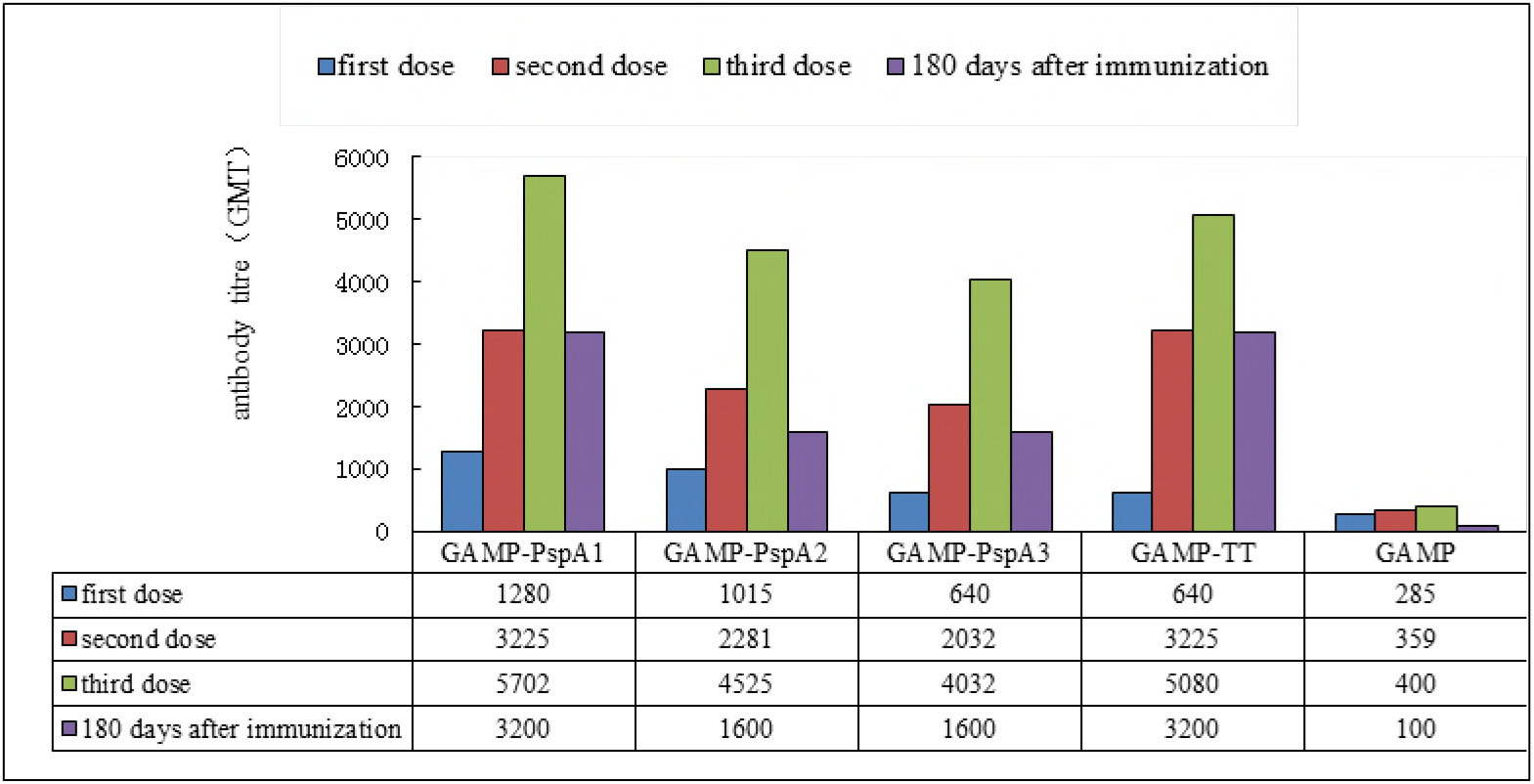
Anti-GAMP antibody titers of the conjugates and un-conjugates after the first, second, third dose and 180 days after the first immunization. The results expressed as the reciprocal of geometric mean titers (GMT).

**Table 2.** Anti-protein IgG antibody levels of the conjugates and un-conjugated proteins after each dose(GMT)

| **Groups** | **First dose** | **Second dose** | **Third dose** |
| --- | --- | --- | --- |
| PspA1 | 45 | 640 | 9051 |
| GAMP-PspA1 | 202 | 1280 | 14368 |
| PspA2 | 113 | 806 | 7352 |
| GAMP-PspA2 | 160 | 2560 | 25600 |
| PspA3 | 403 | 1140 | 51200 |
| GAMP-PspA3 | 202 | 2032 | 7184 |
| TT | 1016 | 10240 | 40637 |
| GAMP-TT | 45 | 113 | 356 |

Note: The results expressed as the reciprocal of geometric mean titers (GMT).

## 3.4 FCM results of the conjugates and un-conjugates

The test mainly detected the condition of immunized mice spleen cells after stimulating with non-specific stimulants in vitro. IFN-γ and IL-4 antibody were used as markers for Th1 and Th2 cell groups. The results of Th1/Th2 ratio were showed in figure3. Compared with un-conjugated GAMP and PBS control, the ratio of Th1/Th2 in groups of polysaccharide-protein conjugates were significantly higher, P < 0.05. The ratio in the three GAMP-PspA conjugates was similar, P>0.05, which were higher than that of in GAMP-TT conjugates, P < 0.05. There were no significant differences between the three un-conjugated PspA groups and the three GAMP-PspA conjugates, P>0.05.

**Figure3.**
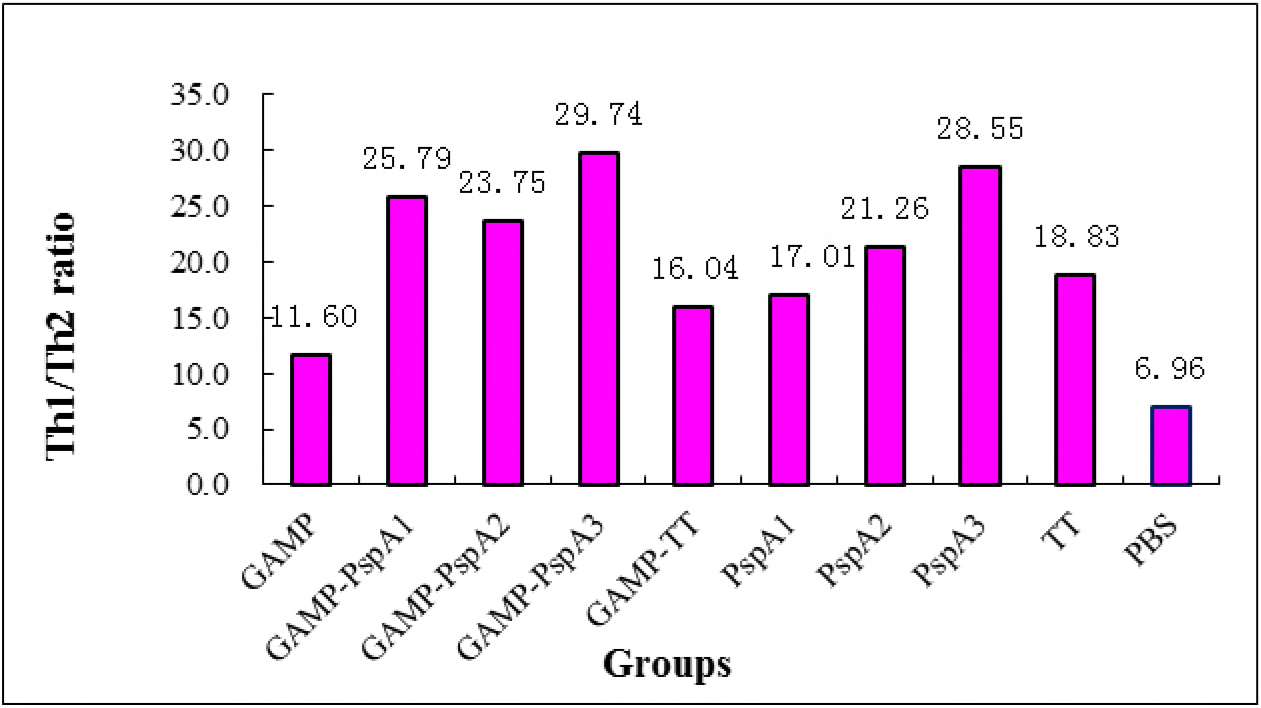
Results of FCM: Th1/Th2 ratio of the conjugates and un-conjugates by FCM in mice 7 days after the third immunization.

## 3.5 Bactericidal activity

Serum bactericidal assay is the most convincingly test, which can directly reflect the potency of meningococcal vaccine. SBA activities were expressed by geometric mean antibody titer. Figure 4 showed the results of test. Anti-GAMP IgG antibody titers in each groups were also showed in the figure, the trend of which were nearly the same with that of SBA results. Pearson correlation was used to analyze the correlation between the results of SBA and antibodies, which indicated the SBA activity is positive correlation with the antibody levels of anti-GAMP, R=0.604, P < 0.05. The polysaccharide-protein conjugates have higher SBA activity compared with that of un-conjugated GAMP, the difference is significantly, P < 0.05.

**Figure4.**
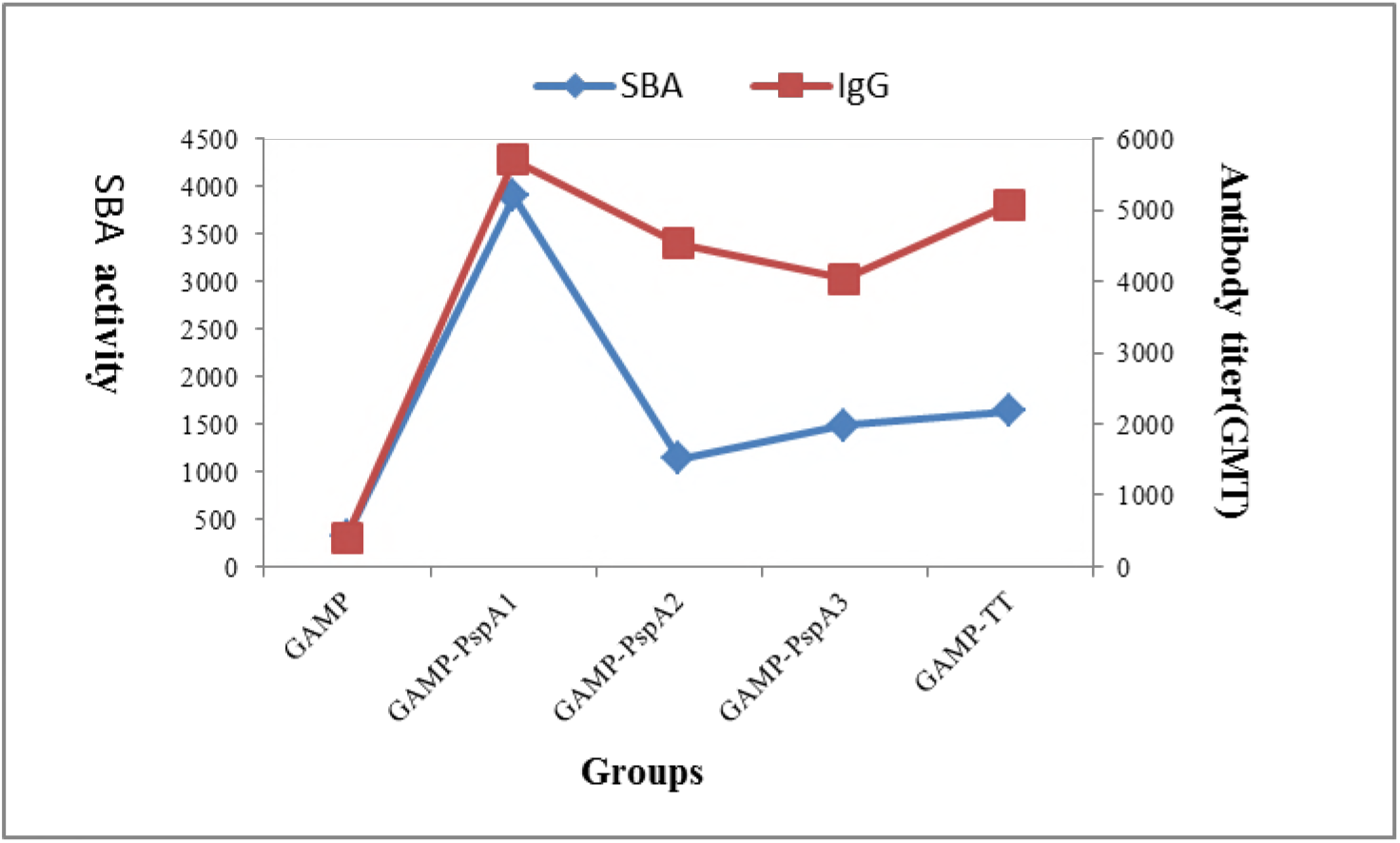
SBA activity and anti-GAMP antibody levels of the conjugates and un-conjugates in mice 7 days after the third immunization. There is a consistent trend and positive correlation between the two (R=0.604, P < 0.05). The antibody results expressed as the reciprocal of geometric mean titers. (GMT).

## 4. Discussion

According to the genome sequences of corresponding protein in GenBank, we adopt the method of chemical synthesis for our targeting genes. This method can omit the bacterial culture process, and can also obtain any kind of bacteria gene sequences that can be found in GenBank. At the same time, codon optimization before the gene sequence synthesis can improve the expression level of target protein. In our study, we get gene sequences of PspA proteins for streptococcus pneumonia strains DBL6A, RX1 and EF3296 from GenBank. The gene sequences were synthesized after being codon optimization, and connected to prokaryotic expression vector PET30a. it is confirmed by sequencing that the correctness of the connection, and the target gene fragment is in line with corresponding sequences in GenBank and the sequence of amino acids have no mutations. The three PspA proteins were highly expressed with soluble formulation, which account for about 30%-40% in total bacterial proteins respectively. After being purification by three steps of chromatography, the purity of them reached 90 percent. The proteins were identified by western blot with human serum of clinically diagnosed pneumonia.

Bacteria capsular polysaccharide is TI antigen, which cannot stimulate T cells to produce cellular immune responses and cannot produce immune memory. Therefore, the researcher combines carrier protein and polysaccharide by chemical method, which changes the TI antigen properties of polysaccharides for TD antigen and solved the above questions (11). PspA was selected as carrier protein of GAMP conjugate for its potential to provide broad cross-reactive protection against S. pneumonia. The ratio of GAMP: PspA is about 0.63, which is similar for families 1 and 2.

The results showed that PspA from both families 1 and 2 were suitable carrier proteins for preparation of GAMP conjugates. Conjugation of PspA to GAMP resulted in a T-cell dependent response to the GAMP as demonstrated by boosting the antibody responses following the second dose and the third dose, which means the induction of immunological memory. Anti-GAMP responses obtained after 3 doses of the GAMP-PspA conjugates were similar to those achieved with GAMP-TT conjugates. Un-conjugated PspA from family 1 was poorly immunogenic compared with that of conjugation of PspA to GAMP, which means the conjugation significantly increased the anti-PspA response. However, the conjugation of PspA from family 2 to GAMP did not increase the level of anti-PspA antibody, which was lower than that of un-conjugated PspA3, P < 0.05, especially after the third dose. The anti-TT response for GAMP-TT is similar to GAMP-PspA3; the immunogenicity of TT was inhibited by the conjugation to GAMP. This may be due to the coverage of protein antigen epitope or due to the protein changes in conformation in the process of combining polysaccharide to protein (12, 13). For TT, another possibility is that the destruction of T cell epitope in the chemical detoxification process, which affecting its immunogenicity (14). The exact cause remains to be confirmed.

In order to evaluate the cellular immune response following the third immunization with conjugates and un-conjugates, FCM methods were used. For Th1/Th2 ratio detected by FCM, compared with un-conjugated GAMP and PBS control, the ratio in groups of conjugates were significantly higher, P < 0.05. The above results indicated that the carrier protein conjugated with GAMP can activate the cellular immune response.

Serum bactericidal assay is mainly detecting the activity of functional antibody in the serum, which is the most direct method in the evaluation of meningococcal polysaccharide vaccine efficacy. It has proved it is associated with the protection of vaccine in the 1860s (15–18),and it has become the gold standard for evaluation of meningococcal vaccine efficacy. The results of SBA showed that the bactericidal activities of the conjugates were significantly stronger than that of un-conjugated GAMP, and the potency of GAMP-PspA1 was higher than that of GAMP-TT, P < 0.05; other PspA conjugates (PspA2, PspA3) had no statistical differences compared with GAMP-TT group, P>0.05.

In conclusion, described here is a procedure for producing a GAMP conjugates using PspAs as carrier protein, and evaluation of the feasibility of PspA proteins. The method of humoral immune response, cellular immune response and SBA assay were used to verify PspA proteins from both families 1 and 2 were suitable carrier for preparation of GAMP conjugates. The research to a certain extent promotes the development of new carrier protein, and it is also the basis for the research of PspA protein as carrier for other kinds of bacterial polysaccharide vaccine. However, there are many areas of study that need to be continued, such as the side effects of PspA and the resistance to pneumococcal infection. It will be studied in detail in subsequent trials.

## Acknowledgments

The authors would like to acknowledge the contributions of Ms. Ruijie Qiao, Mr. Zhiqiang Zhao and Mr. Lin Du, assisting with the test of serum bactericidal assay.

## Abbreviations

PspA: pneumococcal surface protein A
GAMP: group A meningococcal polysaccharide
ELISA: enzyme linked immune sorbent assay
FCM: flow cytometry
SFC: spot forming cells
SBA: serum bactericidal assay
Th: helper T cell
TI: thymus independent
TD: thymus dependent
NIFDC: national institutes for food and drug control
GMT: geometric mean antibody titers
IFN-γ: fibronectin γ
IL-4: interleukin −4
ANOVA: one-way analysis of variance

